# Evidence for dimensional representations and anticipatory dynamics in facial expression perception

**DOI:** 10.64898/2025.12.17.694978

**Authors:** Tyler Roberts, Yong Zhong Liang, Gerald C. Cupchik, Jonathan S. Cant, Adrian Nestor

## Abstract

Expression recognition relies on the ability to distinguish subtle visual differences across a range of facial expressions. Here, we examine the neural representation of dynamic expressions as reflected by electroencephalography (EEG) data in human adults. We find that a wide range of expressions (i.e., 14 emotional and 10 conversational expressions) can be decoded from neural signals, and that their representational structure evinces the classic dimensions of valence and arousal. Critically, we recover, through EEG-based video reconstruction, dynamic representations whose content succeeds in capturing even fine differences across related expressions (e.g., happy-satiated versus schadenfreude). Further, time-resolved decoding reveals anticipatory dynamics that maximize accuracy before the occurrence of an apex expression in the visual stimulus. These results are validated against behavioral data, which yield static reconstructions consistent with their neural counterparts. Thus, our results shed light on the representational basis of expression recognition and serve to recover the dynamic content of visual experience.

## 1. Introduction

Facial expression recognition is a fundamental human ability playing a crucial role in social interactions. Accordingly, extensive work has investigated its neural locus (Fox et al., 2009; Hadj-Bouziane et al., 2008; Haxby et al., 2000; Pitcher et al., 2008; Schwartz et al., 2023; Taubert et al., 2020) and the diagnostic visual information which underpins it (Ekman and Friesen, 1978; Liu et al., 2022; Matsumoto et al., 2008). In contrast, much less is known about the neural processing of facial expressions, such as their representational geometry, and their dynamics.

Recent work has demonstrated the possibility of decoding facial expressions from neural signals using electroencephalography (EEG) (Li et al., 2022; Muukkonen et al., 2020; Smith and Smith, 2019) and functional magnetic resonance imaging (fMRI) (Saarimäki et al., 2022; Said et al., 2010b; Srinivasan et al., 2016). However, neuroimaging work has been constrained by its focus on a handful of basic expressions (Ekman, 1992), such as fear, anger, sadness, disgust, surprise and happiness (but see Saarimäki et al., 2018). Yet, the range of recognizable expressions extends far beyond these six expressions, as shown by recent behavioral and computational work (Cowen and Keltner, 2020; Du et al., 2014; Liu et al., 2022). Decoding a sufficient number of expressions, on par with those recognized by human observers, is crucial for characterizing the geometry of the representational space (Kriegeskorte and Kievit, 2013) underlying expression perception.

A dimensional account, relying on valence and arousal, has been highly influential in the behavioral (Jack, 2013; Mehu and Scherer, 2015; Russell, 1980) and computational (Toisoul et al., 2021) study of emotion. Some results, pointing to the decodability of expression valence from neural data (e.g., in medial prefrontal cortex), have suggested that a dimensional account is also suitable for characterizing neural representations of expression (Skerry and Saxe, 2014). Hence, the present investigation aims to decode a larger set of expressions from neural data in order to derive an expression space and, thus, to assess the validity of a valence-arousal account for neural representations.

Further, we capitalize on the geometry of this space to synthesize the visual appearance of expression representations from the neural signal. While prior work has recovered static representations of facial identity through image reconstruction (Chang and Tsao, 2017; Nestor et al., 2016; VanRullen and Reddy, 2019; Zhan et al., 2019), here we recover dynamic representations of facial expressions through video reconstruction. Motivated by previous efforts with fMRI-based video reconstruction (Nishimoto et al., 2011; Wang et al., 2022), we appeal to the higher temporal resolution of EEG signals to both decode and visualize dynamic information.

Our focus on dynamic expressions, as opposed to static ones, is grounded in their ecological validity, as humans continually change their expressions rather than hold a static one during social interactions (Arsalidou et al., 2011; Trautmann et al., 2009). Moreover, dynamic expressions provide substantially more information and, thus, can support better recognition, especially for subtle displays of emotion (Bould and Morris, 2008; Recio et al., 2013).

Regarding neural resources, dynamic and static expression perception have been found to share common ones, for instance, in the right superior temporal sulcus (rSTS) (Haxby et al., 2000; Johnston et al., 2013; Said et al., 2010b). However, several cortical areas in the posterior STS, MT/V5, and the fusiform gyrus seem to respond preferentially to dynamic expressions (Greening et al., 2018; Johnston et al., 2013). Regarding neural dynamics, static facial expressions can be decoded around 100ms post stimulus onset (Dima et al., 2018) while dynamic expressions may be decoded even earlier (Smith and Smith, 2019). Again though, this prior work relies on a limited number of basic expressions, prompting the need for a more comprehensive investigation.

Therefore, here, we use EEG and behavioral data to address several questions in the field. Specifically, we aim to: (i) decode a multitude of dynamic facial expressions from EEG signals; (ii) characterize the representational geometry of their neural and behavioral representations; (iii) describe the neural dynamics associated with their perception, and (iv) recover the visual content of dynamic expression representations. Thus, our investigation highlights the temporal profile and the visual representations supporting the perception of dynamic facial expressions.

## 2. Methods

### 2.1 Participants

Fifteen healthy adults (9 females; age: 20-31 years) were recruited from the University of Toronto community to participate in the EEG experiment and to complete a behavioral similarity task. However, one participant did not complete data collection and their data was excluded from all analyses. Sample size was similar to those in previous multivariate EEG studies on neural face representations (Nemrodov et al., 2018; Roberts et al., 2019; Smith and Smith, 2019). All participants were right-handed, had normal or corrected-to-normal vision, and reported no history of neurological or visual impairment.

An additional number of 22 White adults (12 female, age: 19-39 years) were recruited through Prolific (www.prolific.com) to provide valence and arousal ratings for our experimental stimuli.

All participants provided informed consent and were financially compensated for participation. The study was approved by the Research Ethics Board at the University of Toronto.

### 2.2 Stimuli

We selected 24 videos displaying expressions for each of two actors (henceforth referred to as ID1 and ID2) from the large MPI Facial Expression Database (Kaulard et al., 2012). A total of 14 expressions were labeled as emotional and 10 as conversational in this database (Table S1). Each stimulus was processed such that, over the course of ten 100ms frames, it started with a same neutral expression (frame 1) and proceeded to an apex expression in the 10th final frame. Where necessary, additional frames were designed by interpolation between existing frames using FantaMorph (v5, Abrosoft) using 91 fiducial points on key facial landmarks (e.g., eyes, lips, and cheeks), resulting in a realistic display of expression change.

Further, one additional stimulus, used only for oddball trials in the EEG experiment, was synthesized from existing stimuli. Specifically, an “angry” expression, missing from the MPI database, was designed for each identity. This was constructed by manually defining fiducial points corresponding to an angry expression and by interpolating multiple frames between a neutral expression and an apex angry expression.

Each video was scaled uniformly, frame by frame, and aligned with roughly the same position of the eyes and the nose, cropped to show only internal features of the face, and normalized with the same mean and root-mean-square (RMS) contrast values for each channel in CIEL*a*b* color space. While a 10Hz rate may lead to unsmooth motion in a video, the alignment of the stimuli, which eliminated any rigid head motion, facilitated relatively smooth displays (Supplementary Videos 1 and 2). The aim of the low rate was to minimize the overlap in the neural signal associated with consecutive frames (see 2.5.1). However, we acknowledge that rigid head movement can provide additional expression information (e.g., the tilt of a head related to disgust) and, also, that a higher rate can provide even more fine-grained temporal information.

### 2.3. Behavioral testing

#### 2.3.1 Face similarity task

Participants completed a similarity judgement task in which they viewed pairs of expressions, separately for each facial identity, and rated their similarity on a scale from 1 to 7 (very dissimilar - very similar). Each trial started with a fixation cross for 200ms followed by pairs of dynamic stimuli displayed simultaneously for 1000ms. The two stimuli subtended a 3.5° x 3.5° angle from 80cm and were placed 1° to the left/right of fixation. Participants could respond only after stimuli completed their animations and the final frame remained on screen until a response was made. Participants judged every pair of expressions, once for each identity, for a total of 552 trials (276 pairs per identity x 2 identities). Data collection was divided into six experimental blocks of equal length (three blocks per facial identity).

In addition, participants completed the Emotional Recognition Index (Scherer and Scherer, 2011) in which they judged the expressions of static faces in a 6-alternative forced-choice task. All participants were able to recognize expressions successfully (mean accuracy = 74.78%, range = 60.00-86.67%, SD = 7.33%) – these results fall well within the normal range for healthy adults (Scherer and Scherer, 2011).

Participants completed the testing within a single 45-minute session. All stimuli were presented against a black background on an LCD monitor with 1920 x 1080 resolution at 60 Hz refresh rate. MATLAB and Psychtoolbox 3.0 (Brainard, 1997; Pelli, 1997) were used to present stimuli, record participant responses, and analyze data.

#### 2.3.2 Valence and arousal ratings

Another group of participants provided valence and arousal ratings for all stimuli. Specifically, participants were presented, in separate blocks, with the final frame (i.e., the most expressive) of the stimuli as well as with the full dynamic expression stimuli. Participants completed two blocks (24 expressions x 2 IDs x 2 repetitions; 96 trials) of still image judgements and one block of dynamic stimulus judgements (24 expressions x 2 IDs; 48 trials).

In each trial, the stimulus was displayed on screen along with two nine-point self-assessment manikins (SAM) (Bradley and Lang, 1994; Sutton et al., 2019) – SAM is a graphic scale that ranges from one end of a spectrum (i.e., frowning face for valence and sleepy face for arousal) to the other (i.e., happy-smiling for valence and excited/wide-eyed for arousal). Participants selected a point on each 9-point scale separately for valence and arousal. Lower scores indicate negative valence and low arousal whereas high scores reflect positive valence and high arousal. Stimuli remained on screen until participants completed both ratings. Each session was completed within 30min.

We rely, in what follows, on valence and arousal ratings obtained for static apex images, which provide more reliable estimates in this context. These ratings were based on a larger number of trials and are likely less cognitively demanding than ratings for full videos (e.g., because of the simpler task). They also provide a natural counterpart to the valence and arousal estimates obtained for our reconstructions, which were evaluated on static apex images rather than reconstructed videos, given the limited temporal smoothness of the frame-by-frame video reconstructions (see supplementary material, Behavioral validation of reconstruction valence and arousal).

### 2.4 EEG experiment: procedure and preprocessing

The experiment was divided across two 3-hour sessions consisting of one practice and 16 experimental blocks. Each block contained 150 trials (i.e., 3 repetitions of 50 dynamic expressions, including oddballs). Each stimulus was displayed for 1000ms followed by an 800 – 900ms fixation cross. Participants pressed a key whenever an oddball expression was displayed.

Data were recorded using a Biosemi ActiveTwo system (Biosemi B.V.). The 64 electrodes were arranged according to the International 10/20 System and the electrode offset was kept below 40mV. Analyses reported below rely on all electrodes – additional analyses targeting occipitotemporal, central, and frontal electrode subsets are reported in the supplementary material (see Expression decoding across different channel groups, for the channel subsets).

The EEG was low-pass filtered using a fifth order sync filter with a half-power cut-off at 204.8Hz and digitized at 512 HZ with 24 bits of resolution. All data were digitally filtered offline (zero-phase 24 dB/octave Butterworth filter) with a bandpass of 0.1 – 40Hz. Then, data were separated into epochs from -100ms to 1600ms. Noisy electrodes were interpolated if necessary (no more than 2 per participant) and epochs were re-referenced to the average reference. Further, artifacts, such as eye-blinks, were removed using Infomax ICA (Delorme et al., 2007).

After removing trials containing artifacts and false alarms, we retained an average of 99% trials across participants – accuracy for oddball detection was at ceiling (range: 97% – 99%). All analyses were carried out using Letswave 6 (Mouraux and Iannetti, 2008) and MATLAB.

### 2.5. Data analyses

#### 2.5.1 Neural decoding

For each participant, we considered three levels of temporal specificity by applying classification to windows of different sizes: (i) temporally-cumulative decoding (TCD) (Nemrodov et al., 2018; Roberts et al., 2019) used large spatiotemporal patterns across a 50-1500 ms interval; (ii) time-resolved decoding (TRD) used consecutive 10 ms-windows between -100 and 1600ms, and (iii) frame-locked decoding (FLD) used 50 – 300ms windows after frame onset. Further, cross-temporal generalization involved training on each 10ms window, and testing on every possible time window.

In more detail, TCD (Nemrodov et al., 2018; Roberts et al., 2019) boosts classification accuracy at the cost of temporal specificity. This is achieved by exploiting spatiotemporal information in a wide temporal interval across an entire array of channels. To this end, here, we concatenated 47,616 features (64 electrodes x 744 time points during a 50-1500 ms window after stimulus onset), separately for each epoch. A wide 1450ms interval was selected, in this respect, to exploit both early and higher-level visual information relevant to face processing (Ghuman et al., 2014; Nemrodov et al., 2018, 2016; Vida et al., 2017).

To estimate the time course of expression processing, TRD was conducted across 10 ms-windows relying on 320-feature patterns (5 consecutive time points x 64 electrodes). The analysis was performed between -100 and 1600ms, by sliding the window one point at a time. Compared to TCD, this analysis provides a finer-grained temporal estimate of discrimination though, by restricting the amount of temporal information used, it does not achieve the same level of discrimination.

Further, FLD assessed expression classification associated with each video frame, at an intermediate level of temporal resolution. Classification was conducted across 50 – 300ms windows time-locked to the onset of each frame, using 8192-dimensional spatiotemporal patterns (128 time points x 64 electrodes). This interval captures a temporally blurred but frame-anchored pattern of activity: it includes early responses to the current frame together with lingering responses to the previous frame and the onset of responses to the subsequent one. Although successive windows overlap, a 250ms-window aims to provide sufficient information for reliable classification at each frame, and for subsequent reconstruction analyses, while limiting signal carry-over across frames. The choice of window length was guided by prior work showing that multivariate EEG patterns within a similar range carry diagnostic facial information (Nemrodov et al., 2018; Roberts et al., 2019). To verify that our findings did not depend critically on this choice, we repeated the analyses with slightly shorter (200 ms) and longer (300 ms) frame-locked windows, which yielded qualitatively similar results (see Results 3.4).

Last, we investigated the transience of information representation over multiple 10ms temporal windows with the aid of cross-temporal generalization analysis by training on each time window, and then testing on every possible time window until 1600ms (Isik et al., 2014). Our cross-temporal decoding analysis resulted in an 822 x 822 matrix in which the diagonal corresponds to the time course estimations mentioned above.

For all types of decoding, epochs corresponding to the same expression display were averaged across repetitions in each block to increase the signal-to-noise ratio (SNR) of the observations. Then, each spatiotemporal feature was separately z-scored across observations and classification was conducted across all possible 552 expression pairs using linear SVM (c=1) and leave-one-block-out cross-validation across 16 blocks.

Next, classification accuracy across participants was assessed using Wilcoxon signed-rank test against permutation-based chance, for TCD, and sign-permutation tests at each time point/window (Xie et al., 2020) for the other types of decoding. Specifically, for the latter, we subtracted chance performance (i.e., 50%) from each participant’s classification accuracy and, then, we multiplied participant-specific data by ±1. For each permutation sample, we recomputed average classification accuracy across participants. The procedure was repeated 10,000 times yielding a permutation-based distribution of the data which allowed us to convert original classification accuracy values into significance values. Multiple comparison correction relied on FDR, for 10ms windows, and Bonferroni for larger (e.g., 250ms) windows.

#### 2.5.2 Expression space

Expression space constructs were derived for each participant by applying metric multidimensional scaling (MDS) to behavior-based and EEG-based estimates of expression similarity. Here, MDS is beneficial both from a theoretical standpoint, as a way to assess and visualize facial expression space, and from a methodological one, as a denoising geometric regularizer prior to image/video reconstruction (see 2.5.3).

For behavior-based spaces, pairwise similarity ratings for dynamic expressions were averaged across repetitions and, then, converted to dissimilarities by computing 8 minus the mean rating, such that larger values corresponded to more dissimilar expressions. These dissimilarities were treated as approximately interval-scaled, a common assumption for 7-point Likert ratings (Norman et al, 2010), allowing the use of classical (Euclidean) MDS. Although similarity ratings can in principle reflect nonlinearities (e.g., due to typicality effects; Townsend et al, 2024), behavioral dissimilarity matrices were well captured by the Euclidean embedding: 11-13 MDS dimensions explained >99% variance of the behavioral data for the two stimulus identities.

For EEG, pairwise decoding accuracies obtained from the relevant temporal windows (TCD or FLD) yielded dissimilarity matrices separately for each identity. Following MDS, the resulting low-dimensional configurations provided a compact approximation of the underlying expression geometry that could be directly compared across modalities and used as the basis for reconstruction (14 dimensions accounting for >80% variance for the two identities).

For visualization and interpretation, we focus on the first two dimensions of the group-average spaces. We further quantify their structure by correlating, for each identity and modality, expression coordinates on these dimensions with valence and arousal ratings of the corresponding static apex images (averaged across participants) and by relating the dimensions across behavioral and EEG spaces. For completeness, we also examine in subsequent analyses how valence and arousal relate to higher-order dimensions (3–10) of the behavioral and EEG expression spaces.

#### 2.5.3 Image and video reconstruction of dynamic expressions

Our procedure for image reconstruction relies on a recent approach capitalizing on the similarity structure of behavioral and EEG data (Nestor et al., 2020). Briefly, behavior-based image reconstruction involves a series of steps, as follows. First, visual features accounting for the structure of expression space were derived separately for each dimension. These features were computed as weighted sums of image stimuli following a strategy akin to reverse correlation (Murray, 2011; Smith et al., 2012). Following conversion to CIEL*a*b, expression stimuli were summed proportionately to their coordinates separately for every dimension. The dimensionality of the space was restricted to 14 dimensions as a trade-off between accounting for most of the variance in the data while minimizing overfitting. The procedure yielded a total of 14 features, or classification images (CIMs), one for each corresponding dimension.

Second, for each dimension, permutation-based CIMs were generated by randomly shuffling the coefficients associated with the stimulus images. Pixel intensities in the true CIM were then compared to the corresponding intensities of pixels in the permutation-based CIMs and only CIMs that contained pixel values significantly different from chance were retained for reconstruction purposes.

Third, the coordinates of the target face were estimated into the existing face space. Importantly, to avoid dependency, the target expression was left out from face space construction and feature derivation by using a leave-one-out approach.

Lastly, a linear combination of significant CIMs, proportional with the coordinates of the target in face space, was added to an average face obtained from all other faces. The outcome of this procedure yields a visual approximation of the appearance of the target for a specific participant.

Neural-based image reconstruction, targeting the apex frame of each stimulus, was conducted in a similar manner relying on TCD results (i.e., relying on a 50-1500ms window after stimulus onset). Further, this procedure was extended to video reconstruction by considering FLD results (i.e., relying on 50-300ms after frame onset).

#### 2.5.4 Evaluation of reconstruction results

Accuracy values were estimated as the relative number of instances for which a reconstructed image/frame was more similar to its target, via an L2 pixelwise distance in CIEL*a*b, than to any other image from a different stimulus (i.e., the final frame for reconstructions of apex expressions or frames with the same number for dynamic reconstructions). Accuracy averaged across participants was compared to chance (50%) via two-tailed one-sample t-tests.

Additionally, a 55-layer neural network (Garcia, 2023) was trained using the Real-World Affective Database (RAF-DB) to represent 18 different expression categories (Li et al., 2017; Li and Deng, 2019). From this, 15,433 images were used for training, with 10% of those images being randomly assigned for validation during training, with a minibatch size of 128 and a learning rate of 0.01. To further validate our reconstructions, we used this network to classify behavioral and temporally cumulative EEG-based reconstructions of apex expressions for each participant. Accuracy was determined as the percentage of instances for which the reconstructed image label matched the label given to the original stimulus image (i.e., the 10th frame of our stimulus video).

## 3. Results

### 3.1. Neural decoding of dynamic facial expressions

To assess the neural discriminability of a range of facial expressions, we recorded EEG signals while participants viewed dynamic expressions (Fig. 1A). Specifically, we used 48 videos of two actors displaying 24 expressions with emotional (e.g., fear) and conversational (e.g., compassion) content. Importantly, the stimuli included variations of basic emotions, such as multiple versions of a happy expression with different semantic content (e.g., happy-satiated, happy-laughing, schadenfreude). Each stimulus started with a neutral expression and evolved to an apex expression (i.e., the most expressive frame) over ten 100ms-frames.

**Figure 1.**
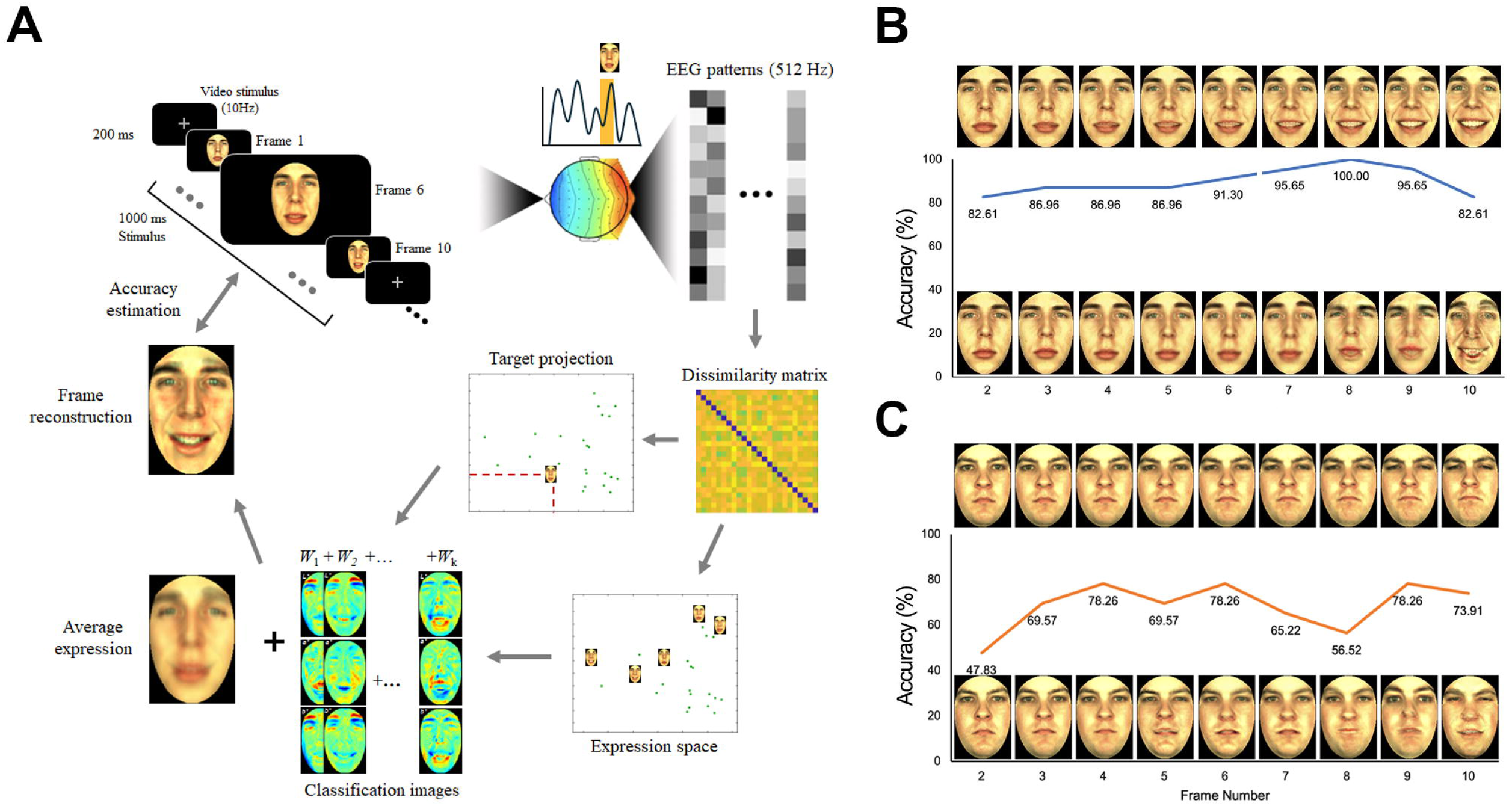
Experimental and analytical procedure along with examples of dynamic expression reconstruction. A) Participants viewed 10-frame, dynamic facial expressions from two actors (facial identities ID1 and ID2). Videos reconstructed, frame by frame, from temporally-constrained EEG patterns aimed to approximate the percepts associated with viewing facial expressions. The procedure relied on neural decoding, expression space estimation, visual feature derivation, image synthesis and accuracy estimation. Representative examples of reconstruction from one participant are shown for (B) ID1 and (C) ID2.

Neural decoding, relying on TCD, assessed pairwise expression discriminability separately for each facial identity (i.e., ID1 and ID2) – for univariate analyses of event-related potentials see Supplemental Results and Fig. S1. Overall, expressions were decoded above chance both for ID1 (mean accuracy across participants: 55.68%; two-tailed Wilcoxon signed-rank test relative to permutation-based chance: V = 103.00, p < 0.001) and for ID2 (mean accuracy: 58.48%; V = 105.00, p < 0.001), with most pairwise decoding accuracies yielding above-chance values for both identities (see Fig. S2 for decoding dissimilarity matrices).

However, these results may be solely due to the discriminability of widely different expressions (e.g., with opposite valence). To examine this possibility, we assessed decoding across expressions with similar meaning and visual appearance. Specifically, we restricted decoding to six expressions under the umbrella category of Happy/Smiling (see Table S1). Again, discrimination was above chance for both ID1 (mean accuracy: 55.36%; V = 97, p = 0.006) and ID2 (mean accuracy: 55.21%; V = 102, p = 0.002).

These results provide evidence for the fine-grained discriminability of dynamic facial expressions from neural signals and, more importantly, they provide a basis for the assessment of a neural-based expression space. Accordingly, next, we estimate such a space based on pairwise discriminability data, we compare it to its behavioral counterpart, and we interpret its dimensions.

### 3.2 Neural and behavior-based expression spaces

To characterize the geometry of expression representations, we derived, for each facial identity, low-dimensional expression spaces from EEG-based estimates of expression discriminability and from behavioral similarity ratings using metric MDS. In both modalities, the resulting spaces showed a meaningful organization: expressions of opposite valence tended to occupy distant regions, conceptually related expressions clustered together, and broadly similar layouts were observed for neural and behavioral spaces (Fig. 2).

**Figure 2.**
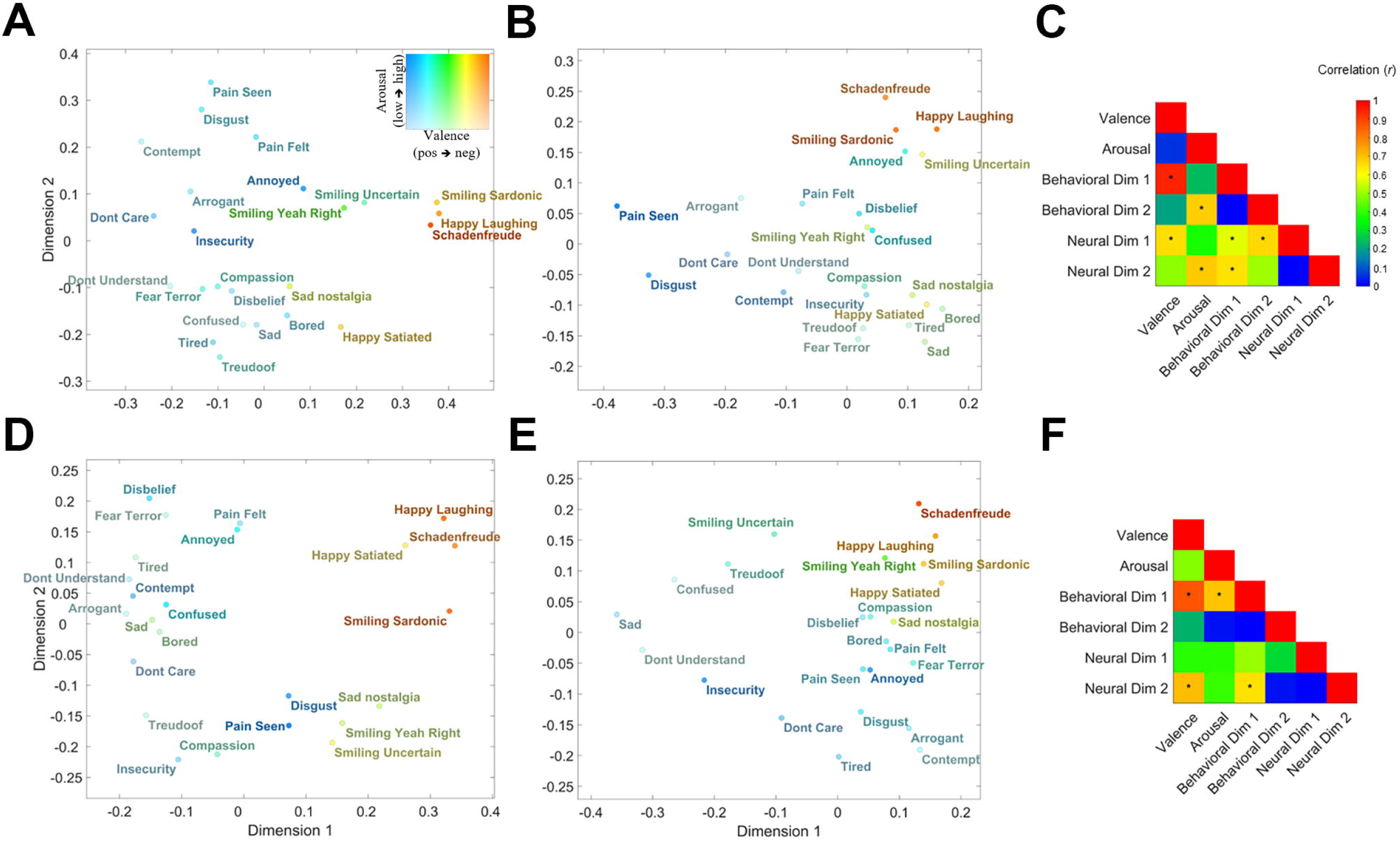
Behavior and EEG-based expression spaces. An approximation of representational spaces derived from A) behavioral and B) neural similarity patterns averaged across participants for ID1. Font hue and brightness indicate valence and arousal, respectively. C) For each data type, behavior and EEG, expression coefficients on dimensions 1 and 2 are correlated across type as well as with expression-specific estimates of valence and arousal, respectively (* *p* < 0.05; Bonferroni-corrected). D-E) Representational spaces and F) correlations are similarly displayed for ID2.

At the same time, expressions from the same broad category (e.g., “happy”) did not collapse into single clusters. For example, ‘happy satiated’ and ‘smiling uncertain’ were displaced from other happy expressions for ID1 (Fig. 2A, B) and ID2 (Fig. 2D, E), respectively, whereas several negative expressions formed tighter subclusters (e.g., ‘bored’, ‘sad’, ‘tired’ for ID1; ‘arrogant’, ‘contempt’ for ID2), consistent with fine-grained representational distinctions among these categories.

Relatedly though, we also note differences in the geometry of the two identities: for instance, ‘annoyed’ and ‘smiling uncertain’ are relatively close for ID1 but further apart for ID2. This likely reflects interactions between identity-related facial appearance and expression (e.g., some physical characteristics associated with a specific facial identity may provide a more compelling depiction of ‘happy’ or ‘sad’). Further, irrespective of objective differences in facial identity, these differences are likely to also reflect how effectively the two actors conveyed particular directed expressions.

A specific hypothesis here concerns the importance of valence and arousal in structuring the geometry of expression spaces. To assess this, we collected valence and arousal ratings for each of our static apex stimuli from a different group of participants (see Fig. S3). We then correlated, for each identity and modality, expression coordinates on the first two dimensions of the neural and behavioral spaces with these ratings, and also correlated these dimensions across data types.

The analysis revealed significant correlations with valence for dimensions of the expression spaces for both identities and both data types (Fig. 2 C, F; p’s < 0.05, Bonferroni-corrected; the correction was applied across 13 of the 15 pairs, as within-modality dimensions are orthogonal by design for both behavioral and EEG spaces). Arousal ratings also correlated significantly with behavioral dimensions across the two identities, but with a neurally-derived dimension only for ID1. Significant cross-modal correlations were also found, particularly for the primary valence-like behavioral dimension. Further correlations between valence and arousal scores, on the one hand, and dimensions 3–10 for either data type and ID did not yield any significant results, indicating that higher-order dimensions encode aspects of expression structure not reducible to these two affective dimensions.

The present findings reveal a meaningful geometry of neural-based representations consistent with a classic model of affect (Russell, 1980). They also demonstrate the correspondence between neural and behavioral representations, especially driven by valence. Last, they make possible a detailed investigation of expression-specific representations and their visualization via image reconstruction as applied to both neural and behavioral data.

### 3.3 Image reconstruction of apex expressions

To uncover the pictorial content of neural representations, we appealed to a recent approach for facial image reconstruction capitalizing on the spatiotemporal structure of EEG data (for review see Nestor et al., 2020). Here, we utilize this technique to assess and visualize representations of static, apex expressions (i.e., frame 10 of our stimuli).

Briefly, a multidimensional expression space was derived from the EEG data of each participant and visual features were derived from of all but one of the expressions in this space (i.e., the reconstruction target). Then, based on the position of the target in the expression space, features were combined into an image reconstruction aiming to recover the appearance of the target expression. Last, reconstruction accuracy was estimated as the proportion of instances for which any given reconstruction was closer to its corresponding stimulus than to any other stimulus. To be clear, here, we consider only the final frame of each of our video stimuli for the purpose of feature derivation and for the estimation of reconstruction accuracy (but see below for the extension of this approach to video reconstruction).

Reconstruction results appear to capture relevant expression information for both facial identities (Figure 3A, B). Accordingly, accuracy was above chance for both ID1 (mean accuracy across participants: 58.44%, two-tailed Wilcoxon signed-rank test compared to 50% chance: V = 103.50, p < 0.001) and ID2 (mean accuracy: 58.45%, V = 103.50, p < 0.001) – see Fig. 3C. A similar approach, which relies on similarity ratings instead of neural discriminability, was applied to derive and assess behavioral-based reconstructions. The outcome of this investigation also evinced above-chance accuracy (ID1 mean accuracy across participants: 75.22%; V = 105.00, p < 0.001; ID2 mean accuracy: 66.56%; V = 105.00, p < 0.001; see Fig. 3C).

**Figure 3.**
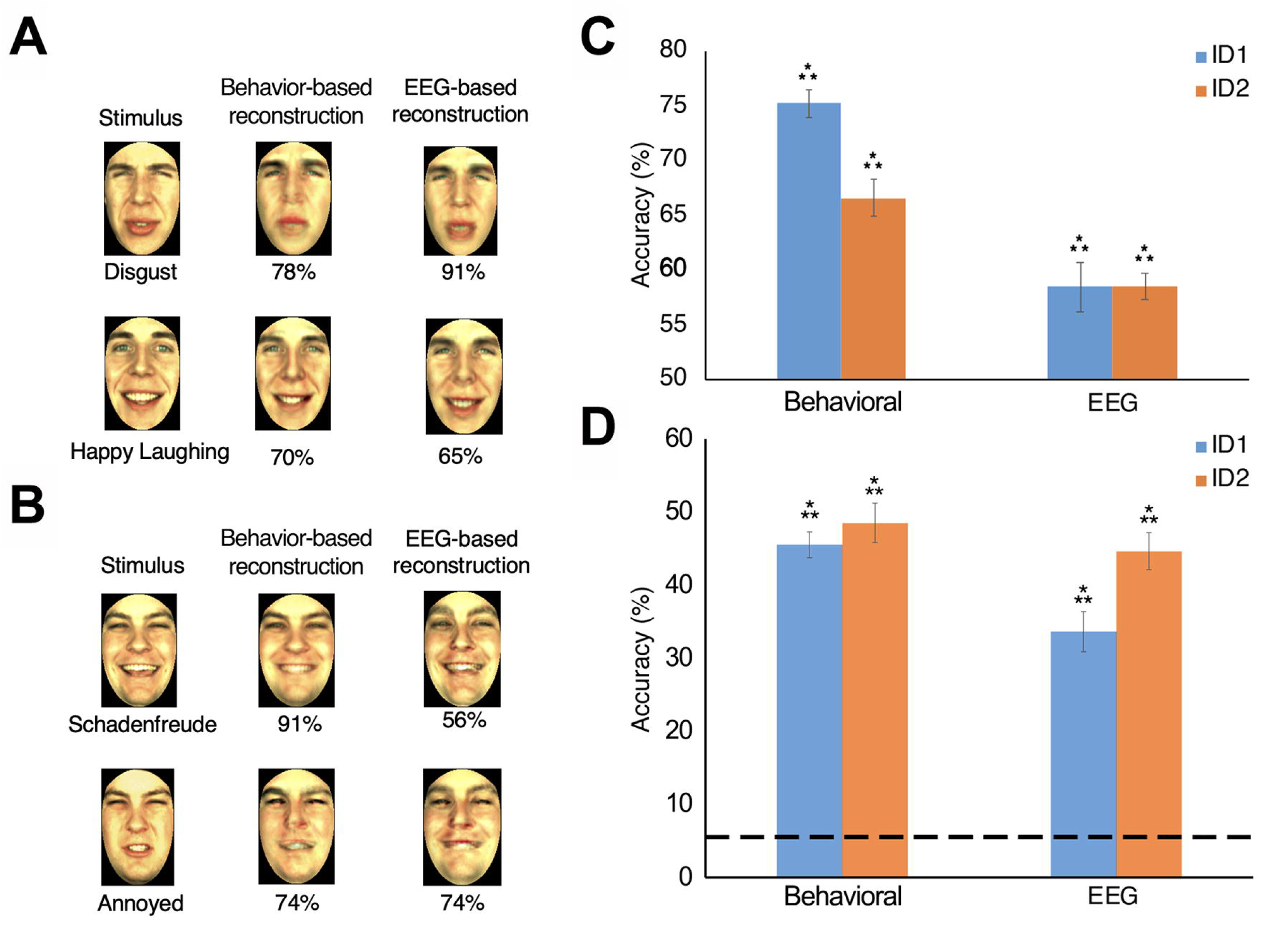
Behavior- and neural-based image reconstructions. Examples of stimuli and corresponding reconstructions from a representative participant for (A) facial identity ID1 and (B) ID2. Reconstructions were based on behavioral similarity ratings and temporally-cumulative EEG decoding corresponding and evaluated relative to the final, most expressive frame of the stimulus videos (numbers indicate image-based accuracy). C) Average reconstruction accuracy based on image-based pixelwise evaluation (chance: 50%). D) Average reconstruction accuracy based on neural network classification (dashed line: 5.56% chance). Error bars indicate ± 1 SE across participants. *** *p* < 0.001.

Further, a two-way repeated-measures ANOVA (2 data types: behavioral and EEG-based; 2 identities: ID1 and ID2) was applied to pixelwise reconstruction accuracies. This revealed significant main effects of data type (F(1, 13) = 40.16, p < 0.001), with higher accuracy for behavioral reconstructions, and identity, with higher accuracy for face identity ID1 (F(1, 13) = 5.06, p = 0.042). The interaction of the two factors was also significant (F(1, 13) = 13.51, p = 0.003), driven by a larger ID1 advantage for behavioral reconstructions relative to EEG.

To provide an alternative assessment of our results, beyond pixelwise similarity, accuracy was re-computed using a deep convolutional neural network (CNN). Specifically, we used a 55-layer CNN (Garcia, 2023) trained with Real-World Affective Database (RAF-DB) images (Li and Deng, 2019). Reconstructions were classified by the network, separately for each participant. Both EEG and behavioral-based reconstructions yielded above-chance accuracy (two-tailed Wilcoxon signed-rank test compared to 5.56% chance, all p’s <0.001) – see Fig. 3D.

Further, a two-way repeated-measures ANOVA (2 data types: behavioral and EEG-based; 2 identities: ID1 and ID2) revealed a significant main effect of data type (F(1, 13) = 12.42, p = 0.004), with higher accuracy for behavioral reconstructions, and a significant main effect of face identity (F(1, 13) = 16.07, p = 0.002), with higher accuracy for ID2. The interaction of the two factors was also significant (F(1, 13) = 14.42, p = 0.002), driven by a larger ID2 advantage for EEG relative to behavioral reconstructions. This pattern of effects mirrors the advantage of behavioral reconstructions observed with pixel-based accuracy estimation. However, it reverses ID1’s advantage found for behavioral reconstructions. This is not a source of conflict though as the two methods of quantifying accuracy prioritize different types of information: lower-level pixel-based versus higher-level visual information diagnostic for expression recognition.

Finally, we asked whether, beyond classification accuracy, the reconstructions preserved the affective qualities of the original expressions. An additional behavioral experiment, in which observers rated the valence and arousal of both behavioral- and EEG-based reconstructions, showed that these ratings closely tracked those for the corresponding apex stimuli (see supplementary material, Behavioral validation of reconstruction valence and arousal).

These findings provide a way to visualize internal representations of facial expressions and to uncover the pictorial information responsible for their successful decoding from EEG signals. Next, we capitalize on the temporal resolution of these signals to characterize the profile of expression decoding and to achieve reconstructions of dynamic representations.

### 3.4 The neural dynamics of expression processing

To estimate the time course of expression decoding, classification was conducted across 10 ms-windows between -100 and 1600ms. ID1 reached above-chance classification at 444ms, peaked at 679ms and exhibited sustained performance between 444 and 1171ms (non-parametric sign permutation tests, FDR-corrected across temporal intervals, q < 0.05; Fig. 4A). In contrast, ID2 reached classification earlier, at 208ms, it peaked later at 810ms, and maintained above-chance classification performance for a longer interval between approximately 362 and 1600ms (Fig. 4A) – for further analyses across different groups of channels see Supplemental Results.

**Figure 4.**
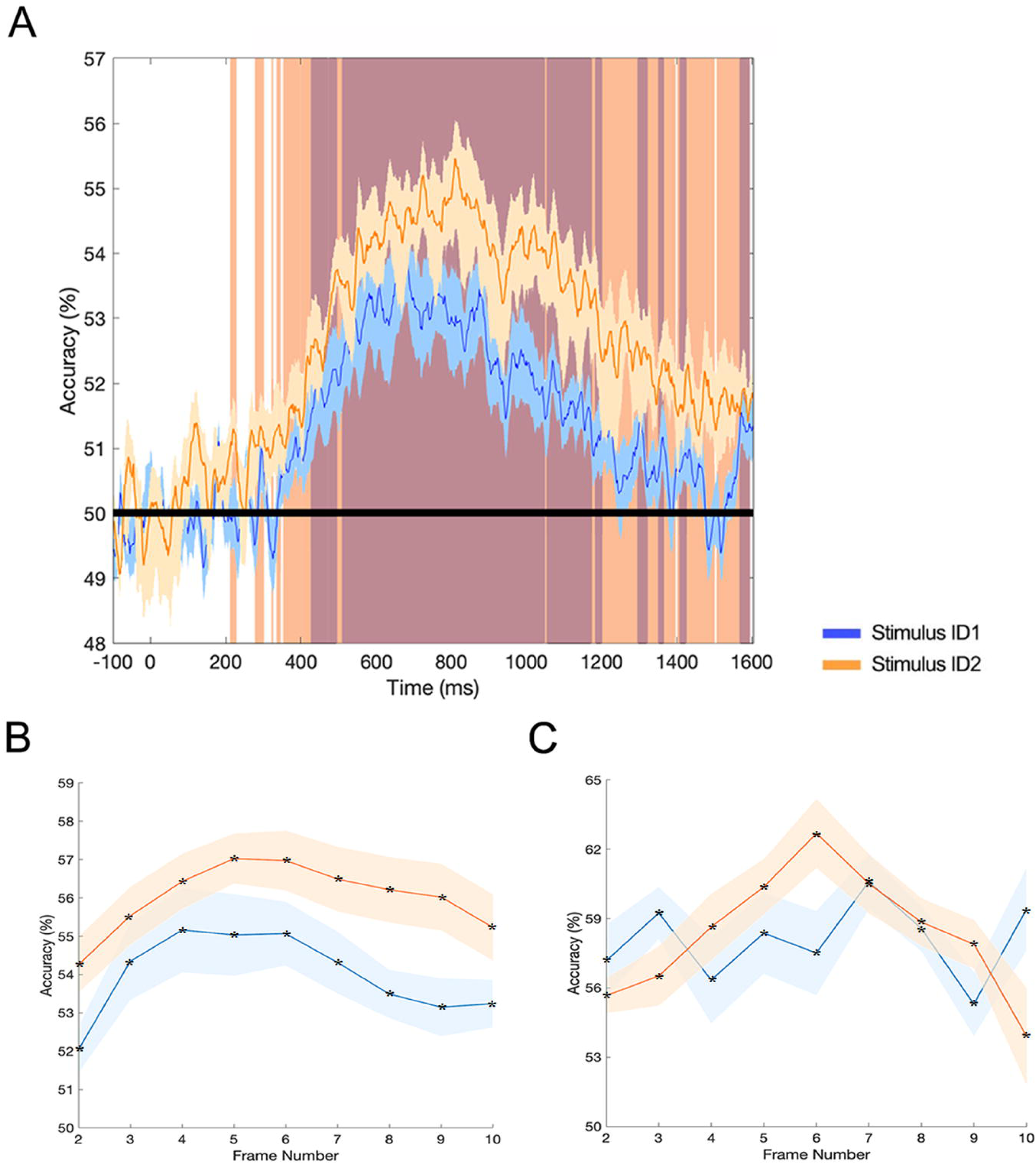
Time course of expression decoding, frame-by-frame classification and reconstruction accuracy of dynamic expressions. A) The time course of expression processing evinced extended intervals of above-chance classification (colored vertical bands; non-parametric sign permutation tests; FDR-correction across time, *q* < 0.05) peaking at 679ms for ID1 and at 810ms for ID2. B) Expression decoding peaked at frame 4 and frame 5 of our stimulus videos for ID1 and ID2, respectively. C) Reconstruction accuracy was carried out for frames 2-10 of each stimulus. Both identities peaked in their reconstruction accuracy at an intermediate frame (7 for ID1 and 6 for ID2). All frame-based expression classification and reconstruction results are significantly above chance for every frame (* Wilcoxon signed-rank test, p < 0.05, Bonferroni corrected). Shaded areas indicate ± 1 SE across participants.

Next, to explore the representation of EEG patterns across time, we carried out a cross-temporal generalizability analysis by conducting pattern classification relying on every possible pairing of time windows for training and testing purposes. Inspection of these results (Fig. 5 and S6) shows evidence for generalization of expression information across time for both face identities. We observe some generalization between 500 and 1100ms for stimulus ID1 and between 300 and 1350ms for stimulus ID2 (q < 0.05).

**Figure 5.**
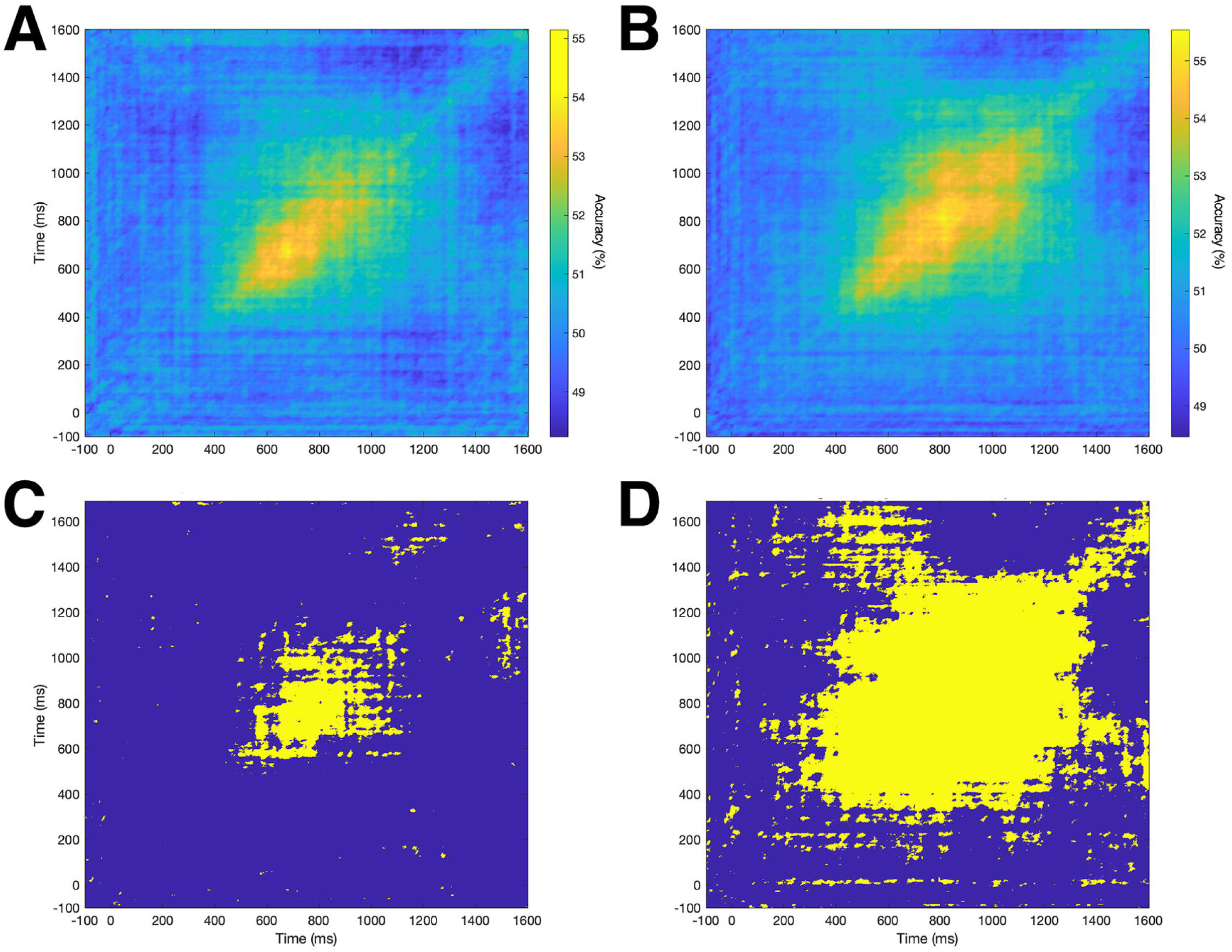
Cross-temporal generalizability of dynamic expression processing. Cross-temporal decoding for facial identity (A) ID1 and (B) ID2 was conducted by training on 10ms intervals (x axis) and by testing on every 10ms interval (y axis) across 64 channels. Time points associated with significant decoding are highlighted (yellow) for (C) ID1 and (D) ID2, respectively (sign permutation test, FDR correction*, q* < 0.05).

While the results above provide a fine-grained assessment of temporal dynamics, their interpretation also has to consider the dynamic nature of the stimuli. Hence, to better relate decoding results to specific visual information in our stimuli, we re-analyzed the data using larger temporal windows associated with different visual frames. Specifically, we conducted classification for windows spanning 50-300ms post frame onset separately for each frame in our stimuli (see Methods for motivation). As shown in Fig. 4B, accuracy is significantly above chance as early as the second frame (i.e., the first frame that diverges from a neutral expression). Then, it remains above chance for all subsequent frames, and it peaks at frame 4 for ID1 (mean accuracy: 55.15%, two-tailed Wilcoxon signed-rank test compared to 50%, V = 105.00, p < 0.001) and frame 5 for ID2 (mean accuracy: 57.03%, V = 105.00, p < 0.001). Interestingly, the frame with the highest decoding accuracy does not correspond to the most expressive frame of the stimulus (i.e., the apex, frame 10), but rather to an intermediate point during the stimulus videos. Comparable frame-locked decoding profiles were obtained when we used 200- and 300-ms windows (Fig. S5A, B), with peak accuracies again occurring at intermediate frames (frames 5–7 for the two identities).

The current findings reveal the neural dynamics of expression processing. Further, they suggest an anticipatory character of these dynamics, which maximize neural discriminability early, ahead of the maximum visual discriminability of the stimuli. This prompts the question whether the present pattern of decoding reflects the higher robustness and accuracy of visual representation at early stages of dynamic stimulus displays. Accordingly, below we assess, via video reconstruction, the content of visual representations associated with each frame and their accuracy.

### 3.5 Neural-based video reconstruction of dynamic expressions

To derive dynamic representations associated with facial expressions, we carried out frame-by-frame reconstructions. To this end, we relied on our decoding results above, based on 250ms temporal windows (i.e., 50-300ms post frame onset), to recover an expression space and its corresponding reconstructions for each frame, expression and participant. Representative reconstructions for ID1 and ID2 are shown in Figure 1B and C, respectively. Reconstructed frames for each identity were further concatenated into videos to better visualize the content of dynamic representations –see Supplemental Videos S1 and S2.

These results evince successful levels of reconstruction for all frames of each stimulus identity (two-tailed Wilcoxon signed-rank test compared to 50% chance; all p’s < 0.01) with the peak for ID1 occurring at frame 7 of the video (average reconstruction accuracy: 60.60%; t(13) = 9.141, p < 0.001) and for ID2 at frame 6 (accuracy: 62.68%; t(13) = 8.488, p < 0.001) – see Fig. 4C. Convergent with our decoding results, this pattern of reconstruction suggests that the accuracy of visual representations is maximized at an early-to-intermediate stage over the course of observing a developing dynamic expression.

Overall, the findings above serve to uncover the dynamic content of expression representations. More generally, they showcase the ability of EEG data to support dynamic reconstructions of neural representations.

## 4. Discussion

The current study investigates the representational content and temporal dynamics underlying the perception of dynamic facial expressions. Our investigation yields several notable findings.

First, we demonstrate that EEG patterns can be used to decode not only a traditional set of basic facial expressions (Muukkonen et al., 2020; Smith and Smith, 2019), but a relatively large and diverse set. Importantly, even within the scope of basic expression categories (e.g., ‘happy’), we can decode different subtypes (e.g., ‘happy satiated’ vs ‘schadenfreude’). Thus, neural decoding can discriminate among expressions with fine differences, both in meaning and physical appearance. This allows a more comprehensive assessment of expression processing and a finer-grained description of neural representations.

Second, we use the structure of our decoding results to recover the representational space of expression perception. Valence and arousal provide a classic framework for the representation of emotions, both elicited and perceived (see Izard, 2007 for review). Here, we present evidence for the validity and utility of this dimensional framework as applied to the neural processing of dynamic expressions (Russell, 1980). Specifically, we show that neural representational spaces are broadly structured by these dimensions and, accordingly, they exhibit significant correspondence with their behavioral counterparts. In other words, valence and arousal emerge as dominant, data-driven axes of the space rather than being imposed a priori, and they explain a substantial portion of the variance in both behavioral and EEG-based similarity patterns. At the same time, successful reconstruction requires higher-order dimensions that do not map straightforwardly onto valence and arousal and reflect additional visual information. Hence, the present results inform a fundamental representational framework in the study of facial expressions.

Third, regarding temporal dynamics, we find that expression decoding reaches above-chance levels relatively early. This is notable given that diagnostic visual information is only available after the first 100ms-frame of the stimulus and that expression-specific cues are subtle at the beginning of a stimulus video compared to its end. Interestingly, decoding peaks for intermediary stimulus frames, and then, steadily declines for the rest of a stimulus. Further, cross-temporal decoding indicates generalizability within a broad interval, roughly between 400ms and 1200ms. Overall, these results suggest that neural processing emphasizes rapidly available, subtle but diagnostic visual information present at the beginning of a stimulus, rather than the abundance of information available later, during an apex display. Beyond the speed advantage conferred by this dynamic profile, another benefit may stem from the ability to clearly distinguish subtle displays of emotion that do not reach an apex within everyday social interactions. Micro expressions, for example, are subtle but recognizable changes in facial expressions which can last less than 250ms (Ekman, 2003; Yan et al., 2013). As micro expressions may indicate the true emotions of a person, there are clear social advantages for picking up on such cues quickly and effectively (Ekman, 2009; Weinberger, 2010).

Fourth, we visualize and evaluate percepts associated with facial expressions by reconstructing their appearance. Specifically, we rely on the structure of representational spaces to recover the visual features and the pictorial content of these percepts. Our results point to successful levels of reconstruction accuracy for both behavioral and neural data, with higher levels for the former, presumably due to the lower SNR inherent in EEG data (Nestor et al., 2020). However, the temporal resolution of EEG data allowed the extension of prior work on image reconstruction (Nemrodov et al., 2018; Roberts et al., 2019) to video reconstruction of dynamic percepts. This leads to a fine-grained visualization of dynamic face representations and opens a path for visualizing dynamic representations more generally. To date, extensive work has aimed to recover the representational content associated with static faces from fMRI (Chang and Tsao, 2017; Nestor et al., 2016; Zhan et al., 2019) and EEG data (Nemrodov et al., 2018; Roberts et al., 2019). Dynamic scene information has also been recovered from fMRI (Nishimoto et al., 2011; Wang et al., 2022). Here, we demonstrate the ability of EEG data to support video reconstruction and, also, we argue for its utility given its superior temporal resolution.

Interestingly, our decoding and reconstruction results are in partial agreement across the two facial identities. Specifically, they evinced broadly similar representational structures and decoding time courses. However, accuracy was lower for one identity (ID1) and an arousal-based dimension for this identity was less apparent in its representational space. These results are not surprising given the interaction of identity and expression processing in face perception (Richoz et al., 2015). However, we note that similarity vectors for the two identities are significantly correlated, suggesting that our methods were reliable in estimating expression spaces across data types and facial identities.

Our results examining different subsets of electrodes yielded significant decoding results across multiple scalp regions, with an advantage for occipitotemporal areas. Given the limited spatial resolution of EEG, these results should be interpreted with caution. While they indicate main reliance on OT information for decoding purposes, overall, they are consistent with the recruitment of a distributed network of cortical areas for processing dynamic facial expressions (Pitcher et al., 2011; Said et al., 2010a). Prior work has demonstrated the role of the pSTS and ventro-temporal cortical areas (Greening et al., 2018; Johnston et al., 2013) along with the motor cortex (Fox et al., 2009) in processing dynamic expressions. More recently, a third stream of visual processing (Pitcher and Ungerleider, 2021) has been proposed to facilitate social perception. This pathway is believed to extend beyond the primary visual areas into the STS and to preferentially process social cues such as gaze direction and dynamic expressions.

We note that our investigation targets the perceptual information underlying expression recognition. However, conceptual information contributes substantially to the neural representation of emotion (Brooks et al., 2019; Cupchik, 2016; Skerry and Saxe, 2015). It is possible that some dimensions of our expression spaces capture higher-level affective and social interpretations. Disentangling these contributions, for example by manipulating orthogonally affective meaning and visual form, represents an important direction for future investigation. More generally, further work is needed to elucidate the integration of these different types of information, the dynamics of this integration, and its significance in the discrimination of subtle differences of facial expression.

An important caveat to our conclusions derives from our use of only two male face identities of the same race. The current study intentionally held identity constant to focus on the richness of expression variation and to simplify the interpretation of the resulting expression spaces, but this necessarily limits the generalizability of our findings. Recent work has successfully applied similar image-based reconstruction approaches to female faces (De et al, 2025), to faces of different races, and to more naturalistic, uncropped face images (Shoura et al, 2025), suggesting that the reconstruction framework itself can scale to more heterogeneous and naturalistic stimulus sets. Hence, future work should not only verify generalizability across a wider range of facial identities but also probe for potential interactions between expression perception and other factors, such as sex, race and visual context.

Overall, the present work provides behavioral and neural evidence for the representational content and the temporal profile of expression processing using a diverse set of dynamic facial expressions. Theoretically, our results speak to: (i) a representational space broadly structured by valence and arousal; (ii) a time course of neural processing evincing sensitivity to subtle but diagnostic, rapidly available visual information, and (iii) a rich pictorial content underlying dynamic representations. Last, methodologically, they demonstrate the capacity of EEG signals to support successful video reconstructions of dynamic face representations.

## Supporting information

Suppementary materials

## Acknowledgments

This research was supported by the Natural Sciences and Engineering Research Council of Canada (J.S.C. and A.N.)

## Author Contributions

J.S.C., G.C.C. and A.N. designed research; T.R., Y.Z.L. and A.N. performed research; T.R., Y.Z.L., J.S.C. and A.N. analyzed data; J.S.C., G.C.C. and A.N. contributed resources and supervision; J.S.C. and A.N. contributed fund acquisition; T.R., Y.Z.L., J.S.C., G.C.C. and A.N. wrote the paper.

## Movie legends

**Movie S1.** Example of dynamic stimulus (*smiling-uncertain* displayed by ID1) and corresponding expression reconstruction from one participant (image-based accuracy is shown for each reconstructed frame). The animation is presented ten times more slowly than the experimental stimulus to facilitate the inspection of individual frames.

**Movie S2.** Example of dynamic stimulus (*smiling-sardonic* displayed by ID2) and corresponding expression reconstruction from one participant (image-based accuracy is shown for each reconstructed frame). The animation is presented ten times more slowly than the experimental stimulus to facilitate the inspection of individual frames.

