## Supplementary material for "Evidence for dimensional representations and anticipatory dynamics in facial expression perception": Suppementary materials

Supplementary methods and results

***Univariate ERP analyses***

Twelve bilateral electrodes located over homologue occipitotemporal (OT) areas (left: P5, P7, P9, PO3, PO7, and O1; right: P6, P8, P10, PO4, PO8, and O2) were used in the ERP analysis – see Fig. S1 inset. These electrodes were selected both because of their relevance to facial expression processing of static (Ashley et al., 2004; Eimer and Holmes, 2002) and dynamic facial expressions (Recio et al., 2013, 2011), as well as because of their ability to support pattern discrimination of different facial properties (Nemrodov et al., 2018; Roberts et al., 2019). Data corresponding to the entire dynamic stimulus display were averaged across electrodes separately for each participant to create a grand average waveform. For univariate analyses, we separately identified the P1, N170, P2, N250, and P4 components.

These components were compared across facial identities and expressions to assess coarse differences in ERP signals evinced by the stimuli used in our study. A two-way ANOVA (2 stimulus IDs x 24 expression stimuli) was conducted separately for each ERP. Our ANOVAs revealed significant main effects of facial identity (ID) for P1 latency (F(1, 624) = 5.40, p = 0.020) and amplitude (F(1, 624) = 1376.32, p < 0.0001) and the latency of N250 (F(1, 624) = 42.85, p < 0.0001). No other significant main effects or interactions were found (all p’s > 0.10). The lack of univariate results associated with expression serves as further motivation for the appeal to multivariate analyses in our study.

***Expression decoding across different channel groups***

To investigate possible differences across different scalp locations three sets of 12 electrodes were selected across OT (see Univariate ERP analyses), central (left: FC1, FC3, FC5, C1, C3, and C5; right: FC2, FC4, FC6, C2, C4, and C6), and frontal areas (left: AF3, AF7, F1, F3, F5, and F7; right: AF4, AF8, F2, F4, F6, and F8) – see Fig. S1 inset. A 3x2 repeated measures ANOVA (3 electrode sets and 2 stimulus identities: ID1 and ID2) investigated potential differences in temporally-cumulative decoding (TCD), across a 50-1500ms window (Fig. S4A). This revealed no effect of stimulus ID (F(1) = 3.40, p = 0.09) and no interaction with electrode set (F(2) = 0.54 , p = 0.59). However, the main effect of ROI was significant (F(2) = 22.71, p < 0.001), as driven by higher accuracy for OT channels relative to both central and frontal ones (both p’s < 0.001).

Further, time-resolved decoding evinced extended intervals of above-chance classification across all channels for ID2 but only across OT channels for ID1 (Fig. S4B-D). We note that this does not contradict the results obtained with ID2 by TCD, which concatenates temporal signals over a larger window and capitalizes on large-dimensional spatiotemporal information to boost classification performance.

Last, we conducted cross-temporal decoding separately for each group of channels. Cross-temporal generalization is apparent for both identities (Fig. S6), though the extent of generalization is sparser for frontal and central channels in the case of ID1. More importantly, we find that OT channels support more robust patterns of generalization for both identities, in agreement with their role in facial information decoding (Nemrodov et al., 2018; Roberts et al., 2019), and that they evince more similarity to the results found using all channels (Fig. 5).

Overall, these results are consistent with main reliance on OT channels for decoding purposes in our main analyses, though additional information is also present at central and frontal scalp regions.

***Image and video reconstruction procedure***

The reconstruction approach aims to recover the visual appearance of each facial expression from its position in an expression space derived from behavioral or EEG data (Fig 1A). For each participant, identity, and modality (behavioral vs EEG), we first constructed a 24×24 dissimilarity matrix describing the relationships among expressions. For behavior, dissimilarities were obtained by converting pairwise similarity ratings into distances (8 - rating). For EEG, dissimilarities were derived from pairwise decoding accuracies, using either temporally-cumulative decoding (TCD) for static reconstructions (50–1500 ms after stimulus onset) or frame-locked decoding (FLD) for dynamic reconstructions (50–300 ms after each frame onset). Metric multidimensional scaling (MDS) was then applied to each matrix to obtain a low-dimensional expression space; we retained 14 dimensions per space, which captured most of the variance (>80% for each participant) while limiting overfitting.

From these spaces we derived dimension-specific visual features, or classification images (CIs), by applying reverse correlation method across stimulus images separately in each CIEL*a*b* color channel. For static reconstructions we used the apex (10th) frame of each video; for dynamic reconstructions the same procedure was applied separately to each of the 10 frames. For a given dimension, we computed a weighted average of all stimulus images, excluding the current reconstruction target, with weights equal to the coordinates of each expression on that dimension. This produces a CI that captures how pixel values covary with the corresponding expression-space axis. To ensure that CIs reflected reliable structure, we generated permutation-based null CIs by randomly shuffling coordinates across expressions and recomputed the weighted averages. Pixel values in the empirical CIs were compared to this null distribution, and only pixels exceeding a predefined threshold were retained (q<0.1, FDR-corrected; 1000 shuffles); dimensions whose CIs contained no significant pixels were dropped from subsequent reconstruction.

Reconstructing a given target expression requires its coordinates in the same space used to derive the CIs, while avoiding circularity by not using the target’s own image to define those features. To achieve this, we used a leave-one-out embedding procedure. For each target, we first constructed an expression space from the dissimilarity matrix with that target removed; this space was used to derive all CIs. We then built a second space from the full matrix including the target. The two spaces were then aligned using a Procrustes transform based on the 23 expressions common to both. Applying the resulting transform to the coordinates of the target in the full space yielded its coordinates in the feature space used for CI derivation, without ever using its image to compute those features.

Static image reconstructions were obtained by linearly combining the significant CIs according to the target’s coordinates and adding this weighted sum to the average of all non-target images. This procedure was applied to apex frames using behavioral expression spaces for behavioral reconstructions and temporally-cumulative EEG expression spaces for neural reconstructions. For dynamic reconstructions, all steps above were repeated independently for each frame: dissimilarity matrices were built from frame-locked EEG decoding, MDS was applied to obtain frame-locked spaces, CIs were derived and thresholded per frame, target coordinates were estimated with the same leave-one-out/Procrustes procedure, and frame reconstructions were formed from the corresponding CIs and average expression for that particular frame. Concatenating the 10 reconstructed frames for each expression yielded a reconstructed video. Because each frame is reconstructed independently from the neural pattern elicited shortly after its onset, temporal continuity is not directly enforced and visible “jumps” can occur between frames, a property we take into account when interpreting and behaviorally validating the reconstructions.

***Behavioral validation of reconstruction valence and arousal***

To provide perceptual validation of the reconstructed expressions, we conducted an additional behavioral experiment in which observers rated the valence and arousal of reconstructions derived from both behavioral and EEG data. Twenty-seven White adults (11 females; age 20–31 years) took part in online testing on Prolific. One participant was excluded due to poor within-participant consistency (negative correlations between ratings for the same stimuli across blocks), leaving 26 participants for analysis. All participants provided informed consent and received monetary compensation.

Because our video-reconstruction procedure operates frame by frame and does not enforce temporal continuity across frames (see Movies S1-S2), reconstructed sequences can contain visible “jumps” that would complicate global judgements of valence and arousal over time. For this reason, we focused on still apex frames, in line with the procedure used previously for rating the original stimuli.

Reconstructed stimuli were generated, for convenience, from dissimilarity matrices averaged across participants in the original experiment, separately for each identity (ID1, ID2) and data type (behavioral vs EEG-based). For each of the four reconstruction sets (2 identities × 2 data types), we considered the apex (10th) frame of each expression, yielding 24 reconstructed images per set.

Participants viewed each set of reconstructions in two separate blocks, with trial order randomized in each block. Each trial presented a single reconstructed apex image along with the same 9-point Self-Assessment Manikin (SAM) scales for valence and arousal used for the original apex ratings. Participants indicated their rating on each scale with a keypress, and the image remained on screen until both responses were recorded. Ratings were averaged across the two blocks for each participant.

For each identity and data type, we then correlated expression-wise mean ratings for reconstructed apex images with the corresponding ratings obtained for the original apex frames (averaged across participants in the original valence/arousal experiment). Separate correlations were computed for valence and arousal, averaged across participants and compared to chance (0) via a Wilcoxon signed-rank test. These analyses yielded above-chance correlations (Bonferroni-corrected) between reconstructions and corresponding original stimuli for both affective dimensions, providing behavioral evidence that the reconstructed images preserve key valence- and arousal-related properties of the underlying expressions (Fig. S7).

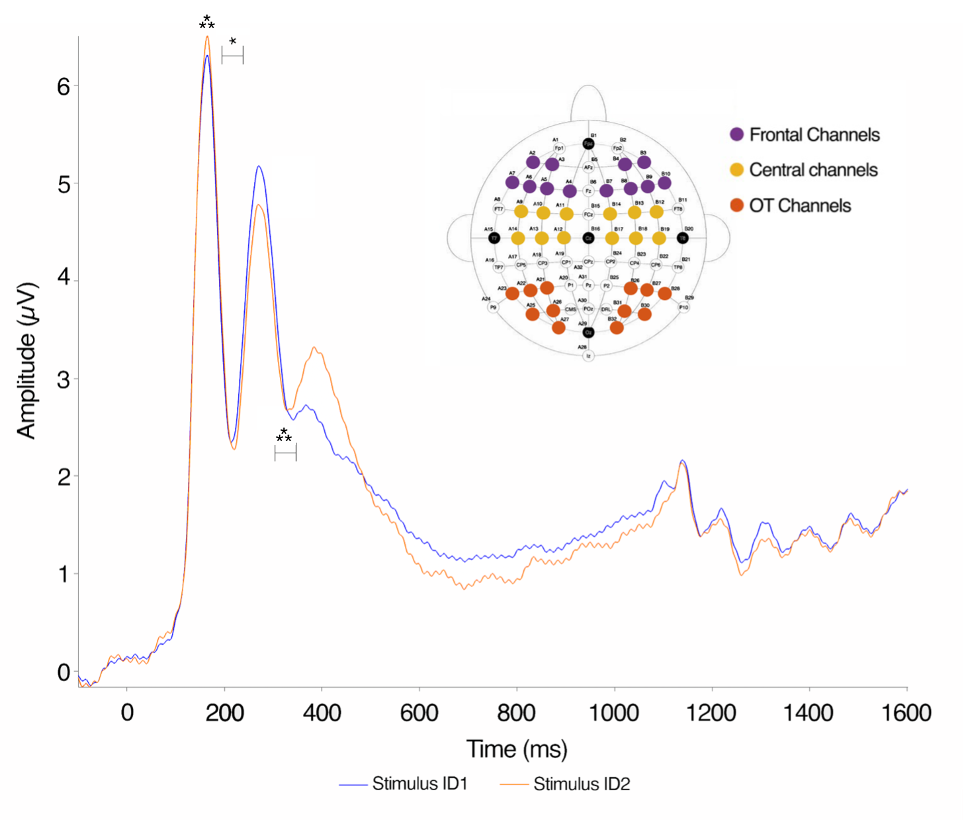

**Figure S1**. **ERP waveforms for dynamic facial expressions from 12 occipitotemporal (OT) electrodes.** A comparison of ERPs elicited by each stimulus identity across 12 bilateral occipitotemporal (OT) electrodes revealed a significantly larger P1 amplitude for ID2, as well as significant latency differences at both P1 and N250. No other amplitude differences reached statistical significance. * *p* < 0.05; *** *p* < 0.001. The inset identifies OT channels (as well as central and frontal channels).

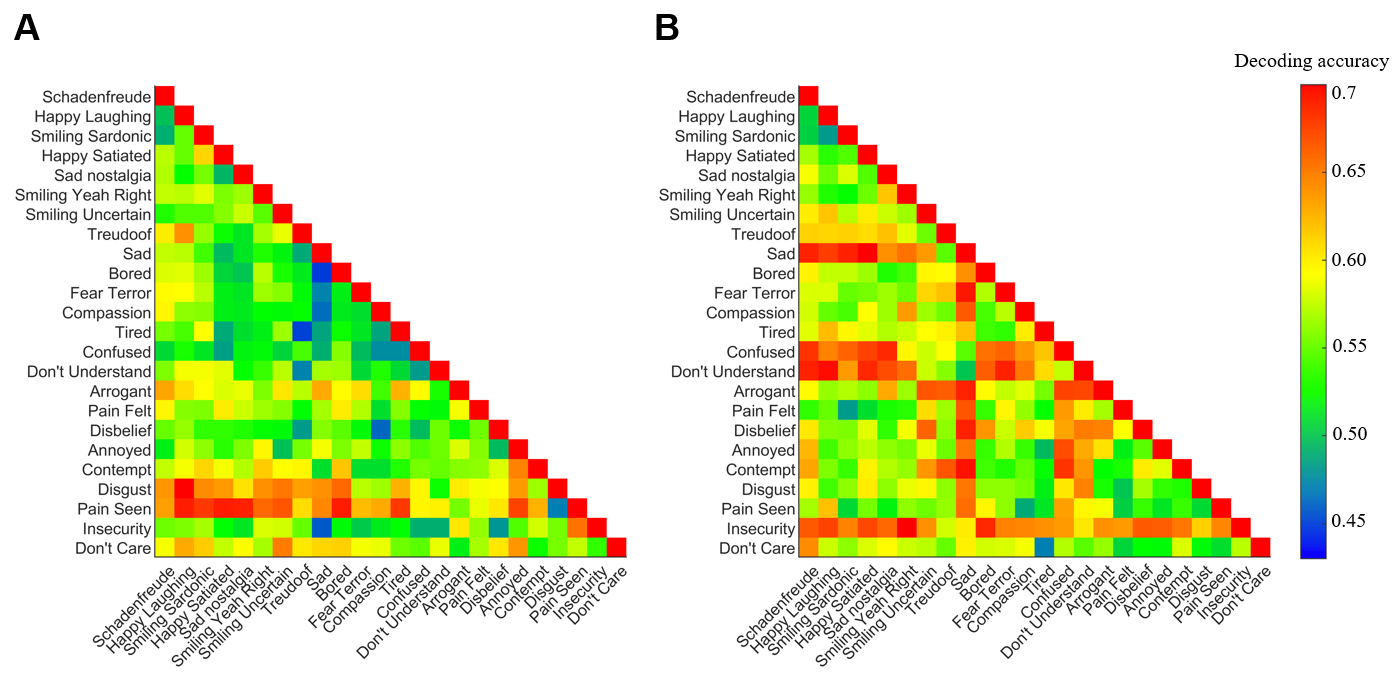

**Figure S2**. **Pairwise expression decoding accuracy.** EEG-based decoding was separately conducted for A) ID1 and B) ID2. Expressions are ordered by valence ratings, averaged across the two stimulus identities to facilitate their comparison. A majority of pairs yield above-chance decoding accuracy.

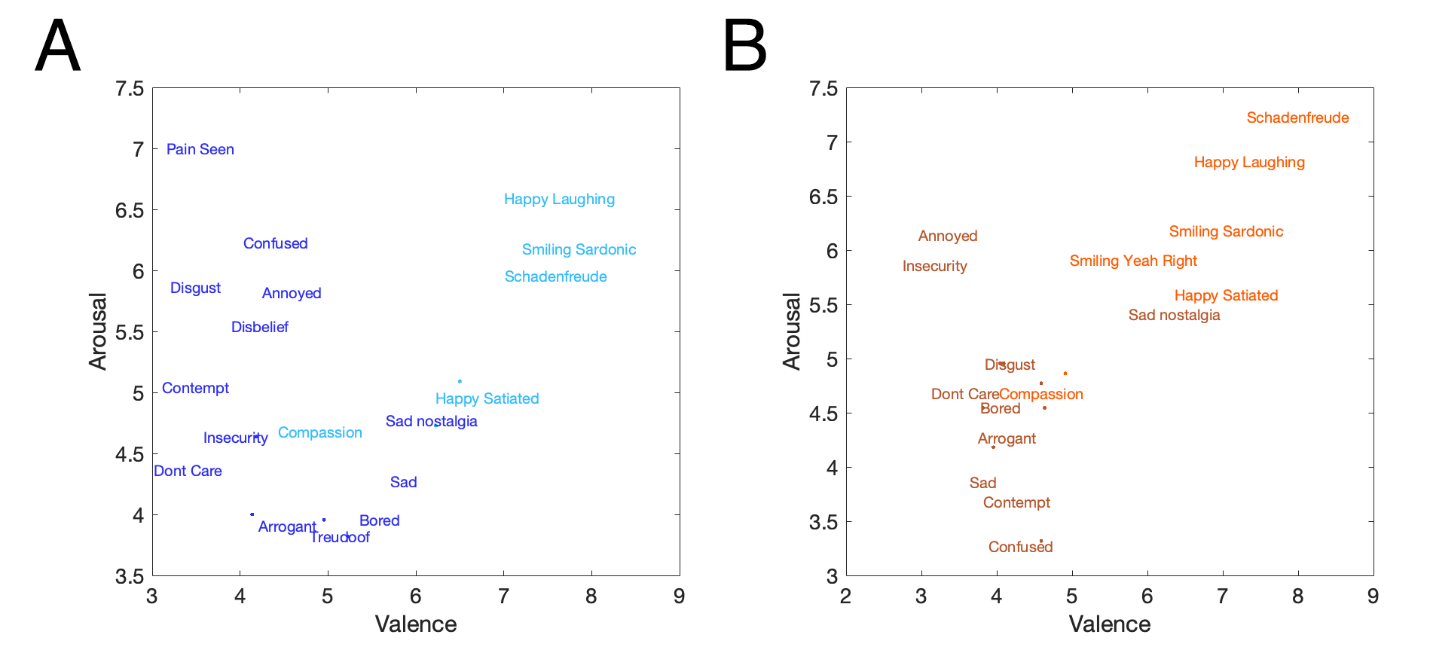

**Figure S3.** **Valence and arousal scores for expression stimuli.** Average scores of valence and arousal judgements across participants for (A) ID1 and (B) ID2. Darker and lighter fonts indicate expressions with negative and positive valence, respectively.

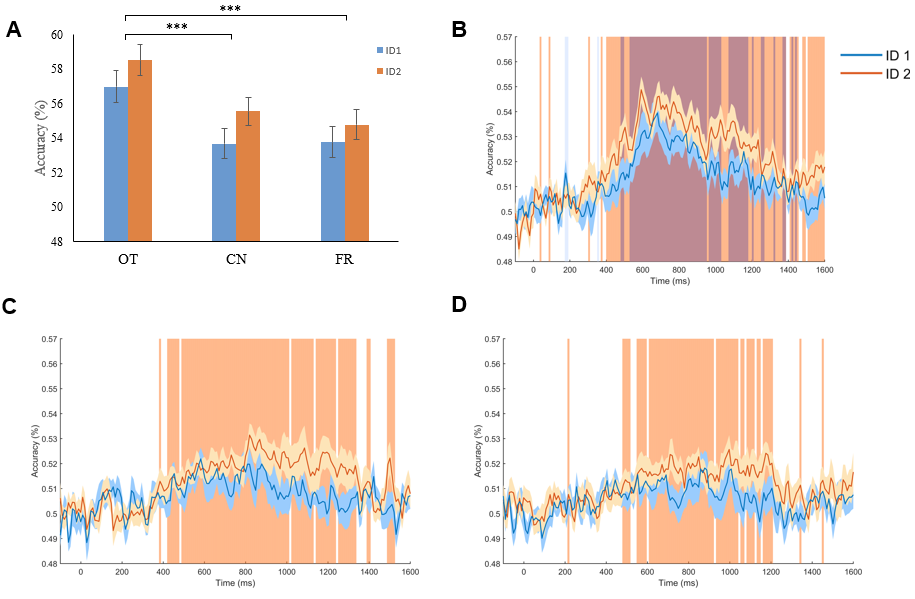

**Figure S4.** **Decoding accuracy for different channel sets.** A) Temporally-cumulative decoding yielded above-chance accuracy for all channels sets and for both stimulus identities as well as an advantage for occipitotemporal (OT) channels over central (CN) and frontal (FR) ones (*** p<.001, Bonferroni-corrected). Time-resolved decoding was separately conducted for B) OT, C) CN and D) FR channels. The time course evinced extended intervals of above-chance classification across all channels for ID2 but only across OT channels for ID1. Time points associated with significant decoding are highlighted for each face identity (sign permutation test, FDR correction, q < 0.05). Shaded areas indicate ± 1 SE across participants.

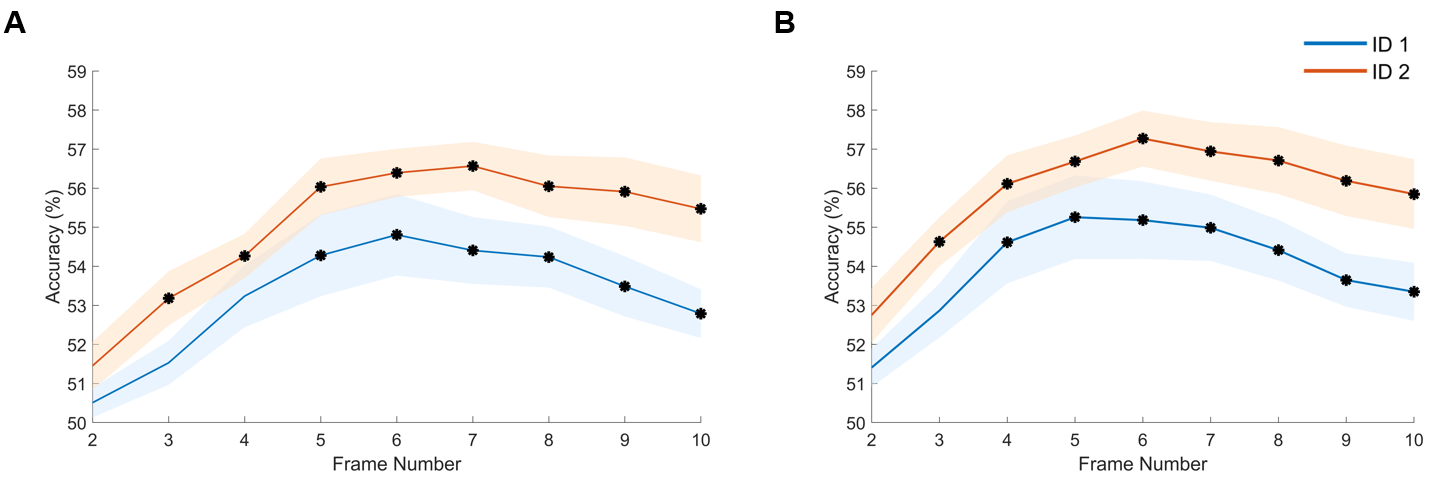

**Figure S5.** **Frame-locked expression decoding.** Decoding was performed using A) 50-250 ms and B) 50-350ms windows after frame onset for frames 2-10 of each stimulus. (sign permutation test, FDR correction*,* *q* < 0.05). Both identities peaked in their reconstruction accuracy at an intermediate frame. Overall, decoding accuracy increases with the size of the window while limiting temporal and frame specificity. Shaded areas indicate ± 1 SE across participants (* p<.05; Wilcoxon signed-rank test, Bonferroni correction across frames).

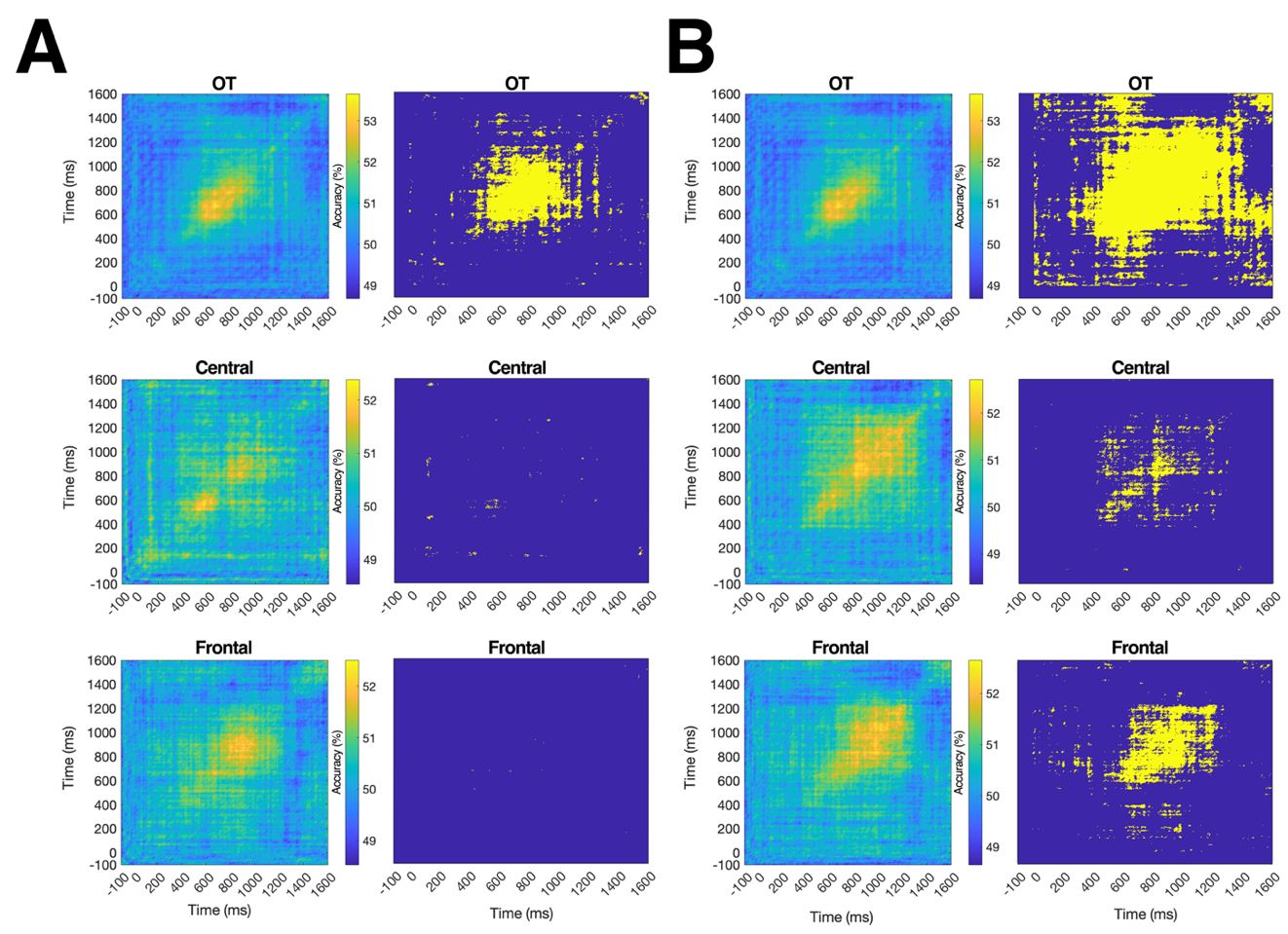
**Figure S6.** **Cross-temporal generalizability of dynamic expression processing for different channel sets.** Cross-temporal decoding for (A) ID1 and (B) ID2 was conducted by training on 10ms intervals (x axis) and by testing on every 10ms interval (y axis) for different groups of electrodes (occipitotemporal, central, and frontal). Time points associated with significant decoding are highlighted (yellow) for each face identity next to their decoding heatmaps (sign permutation test, FDR correction*,* *q* < 0.05).

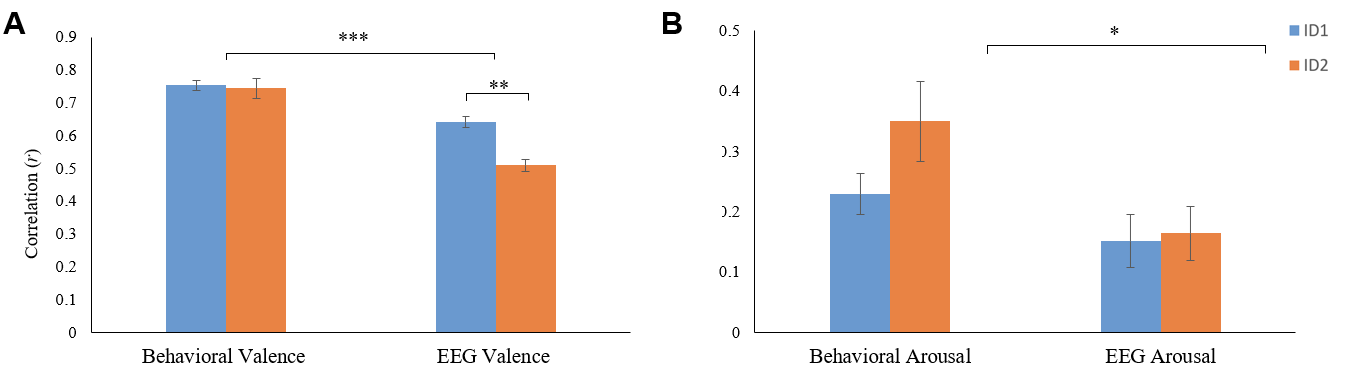

**Figure S7. Behavioral assessment of image (apex) reconstructions.** A) Valence and B) arousal ratings were consistent across stimuli and their corresponding reconstructions from behavioral and EEG data. Behavioral data yielded higher correlations than EEG ones (***, p<.001; ** <.01; * p<.05). Error bars indicate ± 1 SE across participants.

**Table S1. Expression labels and their category.** A total of 24 dynamic expressions were selected from the MPI database (Kaulard et al., 2012), consisting of 14 emotional expressions and 10 conversational expressions.

| **Emotional Expressions** | **Conversational Expressions** |
| --- | --- |
| Arrogant | Annoyed |
| Contempt | Bored |
| Disgust | Compassion |
| Fear | Confused |
| Happy Laughing | Disbelief |
| Happy Satiated | Don't Care |
| Pain Felt | Don't Understand |
| Pain Seen | Insecurity |
| Sad | Tired |
| Sad Nostalgia | Treudoof (‘doe-eyed’) |
| Schadenfreude |  |
| Smiling Sardonic |  |
| Smiling Uncertain |  |
| Smiling Yeah Right |  |
